# Wearable-ome meets epigenome: A novel approach to measuring biological age with wearable devices

**DOI:** 10.1101/2023.04.11.536462

**Authors:** Cameron Sugden, Franco B du Preez, Laurence R Olivier, Armin Deffur

## Abstract

Aging is an inevitable process of cellular and physiological decline. These markers of age can be measured on the molecular and functional level. Wearable devices offer a non-invasive continuous measure of physiological and behavioural features and how they pertain to aging. Wearable data can be used to extrapolate information derived from epigenetic biological age predictions and its underlying biology. LifeQ-enabled wearable devices were worn for 40 days to harvest data on 48 human participants. Thereafter blood was drawn and methylation levels determined using the Illumina EPIC array. Multiple epigenetic clock ages were calculated and compared with wearable features. Activity minutes correlated with VO_2_ max (p = 0.003), subendocardial viability ratio (SEVR, p < 0.01), blood pressure index (BPI, p = 0.02), resting heart rate (RHR, p < 0.01) and heart outflow (HO, p < 0.01). Sedentary time correlated with RHR (p < 0.01), VO_2_ max (p = 0.01), SEVR (p = 0.04), and HO (p = 0.04). VO_2_ max, SEVR, small artery resistance (SAR), BPI and large artery stiffness index (LASI) correlated with multiple epigenetic age clock outputs and chronological age but were most strongly correlated with PCPhenoAge. VO_2_ max, (p = 0.04) RHR (p < 0.01) and LASI (p = 0.04) were significantly correlated with PCPhenoAge acceleration. Weighted gene correlation network analysis (WGCNA) of the differentially methylated positions of PCPhenoAge acceleration was used to construct modules, identifying 3 modules correlating with wearable features. Behavioural features impact physiological state, measured by the wearable, which are associated with epigenetic age and age acceleration. Signal from the underlying biology of age acceleration can be picked up by the wearable, presenting a case that wearable devices can capture portions of biological aging.

## Introduction

### The concept of biological age

Aging is the time-dependent process of cellular and physiological decline of an organism, ultimately, leading to death^1^. Age is accompanied by an increased risk for age-related diseases such as cardiovascular disease, cancers, neurodegenerative diseases,^2^ and death. The average human life expectancy has increased over the last decade, and projections estimate that by the year 2050 at least 25% of the developed world population will be over the age of 65^3^; however, healthspan, the portion of life free from chronic disease, is not increasing in the same manner^4^. Understanding the molecular and physiological mechanisms of aging, how to track them, and how to influence the underlying processes and pathways may provide an opportunity to delay the onset of chronic disease and prolong both healthspan and lifespan. This hypothesis introduces the idea that aging is a dynamic process and simply utilizing chronological age as a predictor of all-cause mortality (ACM) and disease onset is insufficient. Instead, measuring the organismal state of function, biological age, may provide greater insight.

### Hallmarks of aging

Biological aging is paired with a physiological state of increased risk of disease and decreased function^2^. To better understand the molecular footprint of aging, Lopez-Otin, *et al.* identified the hallmarks of aging^5^, categorized as follows: *primary hallmarks* (the cause of damage), including genomic instability, telomere shortening, loss of proteostasis, and epigenetic modifications; *antagonistic hallmarks* (the response to damage), including dysregulated nutrient sensing, mitochondrial dysfunction, and cellular senescence; and *integrative hallmarks* (the consequences of the phenotype), including stem cell exhaustion and altered cellular communication^1^. The hallmarks provide a landscape to understand the intricate and complex networks present in aging. For example, the aging phenotype could have a direct link to the specific disease, such as mitochondrial dysfunction and neurodegenerative diseases^6^. Alternatively, systemic changes could be an indirect cause of aging-related diseases, such as an inflammatory response to senescent cells elevating levels of chronic inflammation leading to detrimental downstream effects^7^. As mentioned above, alterations of the epigenome are considered a primary hallmark of aging^1^. One of these modifications, in mammals, is DNA methylation (DNAm) of the cytosine residue usually found at cytosine-guanine dinucleotides (CpGs)^8^. As aging progresses, epigenetic modifications occur on multiple levels. Firstly, in a predictable fashion, termed differential methylation, where methylation status of specific CpG sites change predictably, and secondly in a variable fashion, termed variable methylation, where methylation status of sites vary as aging progresses^9^. Epigenetic clocks are molecular aging biomarkers, which are designed to determine biological age based on measuring the epigenetic modifications that accompany aging.

### Epigenetic clocks

Multiple clocks have been developed, most commonly via a penalized-regression model^10^. Clocks have been trained to predict chronological age (such as Horvath^11^ and Hannum^12^), mortality (such as Lu’s GrimAge^13^), and the risk of age-related disease (such as Levine’s PhenoAge^14^) which was designed to predict “phenotypic age” - a blood biomarker-based measurement capturing morbidity and mortality^15^. DNAm age has been shown to be predictive of ACM, healthspan, and physical function^14^. An additional layer of biological context is revealed when addressing the biological age acceleration, defined as the residual of the linear model of biological age regressed onto chronological age^11^. This error is proposed to be biologically meaningful. An increased age acceleration is associated with decreased lung function^16^, cardiovascular disease^17,18^ and risk factors^19^, neurodegenerative diseases^20,21^, cancer^22^, and mental illness^23,24^. Age acceleration has also been associated with lifestyle factors, negatively correlating with exercise^25^ and positively with sleep disorders^26,27^ and smoking status^25^. On a molecular level, epigenetic acceleration has been associated with key pathways involved in cellular homeostasis, inflammation, cell division, and metabolism, to name a few^14^. A question which comes to mind is which of the many DNAm epigenetic clocks is the “gold standard” clock, if any? A drawback on using these clocks as molecular biomarkers of aging is the technical variance introduced on the individual level^28^. Repetitive measures can result in large variance, which makes the clock unreliable as a biomarker of aging. To address this problem, a new method of calculating the clocks was proposed by retraining the clocks using a principal component analysis (PCA) method^29^, significantly decreasing the variation of repeated testing. Epigenetic clocks also present friction in the invasive and expensive nature of the biomarker, although recent work suggests efforts to dramatically decrease the costs of analysis^30^. Due to the challenges set out above, repetitive measures of epigenetic age to analyse an individual’s age rate or assess the potential impact of lifestyle changes may prove to be difficult. An accurate, non-invasive, continuous estimate of biological age which can provide additional context is lacking. Changes in aging occur on the molecular, cellular, structural, and functional level. Linking these levels of the aging phenotype will aid in understanding the phenotype and how to monitor it.

### Functional biomarkers of aging

Functional biomarkers of aging are biomarkers which provide insight to organ or system function with regards to aging. Grip strength, a proxy for muscular function, for example, has been shown to decrease with age^31^. VO_2_ max, the maximum capacity of the muscle to utilize oxygen during exercise, can be used as a measure of maximal cardiovascular fitness^32^ and has been shown to decline with age, which may be due to decreased cardiac output with age^33^. Structural changes to blood vessels, measured by changes in pulse waveform (PWF) features, have been shown to change with age^34^ and overall vascular health status^35^. Linking functional biomarkers to molecular biomarkers will generate a clearer understanding of the cause-and-effect relationship and be able to identify potential areas where, independently, the two classes of biomarkers capture different aspects of the aging process.

### Wearable devices

Wearable devices are strong candidates to measure the functional biomarkers of aging in a frictionless reliable way. Photoplethysmography (PPG) is a non-invasive method to measure changes of blood volume in tissue. Typically, a LED is shone onto the skin and is absorbed by the blood and tissue, and the remaining light is reflected and measured by a photodiode^36^. As the heart beats, blood volume in the tissue changes, causing the corresponding light which is reflected to change. Once this signal is processed, biological data, such as the PWF, heart rate, heart rate variability and blood oxygen levels can be extrapolated^37^. In addition to PPG, an accelerometer captures sedentary and activity data and has been used to capture such data from large population cohorts^38^. LifeQ Inc., a wearable device biometric company, has the capability to produce a wide range of wearable-based biometric features. These features can be categorized as behavioural or physiological. In line with several omics fields describing a comprehensive collection of information within a particular domain of biology, e.g. biome, genome and transcriptome, we define the wearable-ome as the set of behavioural and physiological biometrics that state-of-the-art technology can monitor longitudinally in the general population. Behavioural features refer to features which are directly impacted by an individual’s actions or habits, such as the frequency and duration of an exercise session. A physiological feature refers to a measure of the individual’s physical makeup or function, such as the individual’s resting heart rate. A description of behavioural and physiological features can be viewed in Table 1. Mapping the wearable features to a molecular measure of biological age could aid in the tracking of biological age and how it can be impacted by behavioural adaptations. Additionally, wearable devices provide the ability to visualize continuous longitudinal data, aiding in the tracking of the aging phenotype and the generation of personalized behavioural interventions to positively impact aging.

**Table 1:** LifeQ’s wearable features (subset used in study).

| Wearable feature | Feature category | Definition |
| --- | --- | --- |
| Resting heart rate (RHR) | Physiological | Frequency of heartbeats measured during a state of rest. |
| VO<sub>2</sub> max | Physiological | Maximal capacity of oxygen uptake and utilization by the muscle. |
| Normalized crest time | Physiological | Indication of the contraction strength of the heart muscle. The stronger the cardiac muscle is, the faster the left ventricle can inject stroke volume into the aorta. |
| Subendocardial viability ratio (SEVR) | Physiological | Indication of the amount of oxygenated blood volume or blood supply to coronary arteries. A high SEVR indicates good blood flow to the heart, while a low SEVR indicates poor blood flow. |
| Large artery stiffness index (LASI) | Physiological | Indication of the stiffness of the aorta and the wave travel speed in blood. Higher LASI is indicative of the higher wave travel speed and, correspondingly, to higher artery stiffness. |
| Ejection duration / Heart outflow (HO) | Physiological | Representation of the blood pumping strength of the heart. A healthy HO value should not be too large or too small. |
| Normalized pulse width / Small artery resistance (SAR) | Physiological | Measurement of the resistance to blood flow in the small arteries. SAR is related to arterial stiffness of the peripheral arteries. Smaller SAR indicates healthier peripheral arteries. |
| B:A ratio / Blood pressure index (BPI) | Physiological | Measurement of the blood pressure. BPI value is proportional to the average blood pressure. |
| Activity time | Behavioural | Activity time per day in minutes, stratified by intensity, light, moderate and vigorous. |
| Sedentary hours | Behavioural | Duration in hours per day in which a state of inactive behaviour was measured. |
| Total sleep time | Behavioural | Duration from sleep onset to wake up, excluding awakenings. |
| REM sleep percentage | Behavioural | Percentage of time for which REM sleep was measured throughout the night. |
| Deep sleep percentage | Behavioural | Percentage of time for which deep sleep was measured throughout the night. |

### Aims and objectives

This study aims to determine to what extent wearable data can be used to extrapolate information derived from CpG methylation and biological age predictions. This was achieved by recruiting individuals of various ages and sampling wearable and blood data. The hypothesis is that there will be an overlap of information provided by alterations in DNAm and wearable features, both physiological and behavioural.

## Methods

### Participants and recruitment

Ethical approval for this study was provided by Pharma-Ethics (211224438). Following the approval, A convenience sample of participants was recruited by a messaging campaign (SMS, word of mouth) aimed at the general population resident in the study catchment area, without regard to health status. The recruitment methods are illustrated in Figure 1 below.

**Figure 1:**
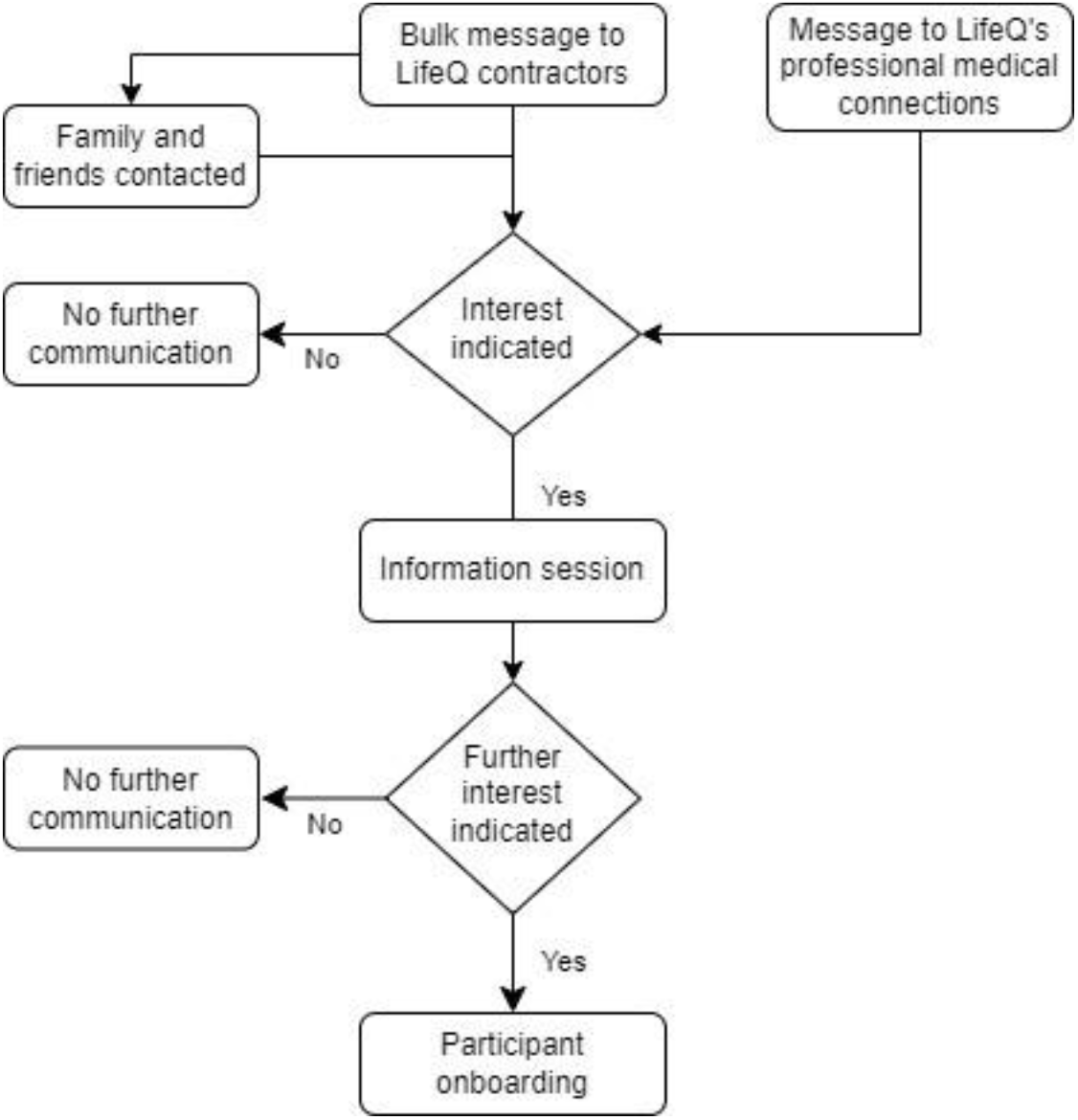
Participant recruitment strategy.

### Study design

Participants were supplied with a Motorola Moto 360 wearable and instructed in its use. The Moto 360 is enabled with LifeQ’s biometrics library utilizing 25Hz PPG and accelerometer sensor signals to produce advanced biometrics. This process involved anonymising their data, connecting the device to LifeQ’s cloud environment, and ensuring data was flowing from the device to the cloud. Participants were encouraged to wear the device for as long as possible during the day, especially during sleep and exercise. Following this onboarding process, the participants completed 40 days of free-living. At the end of the 40-day period, blood was drawn from the participants, and the data collection was terminated (Figure 2).

**Figure 2:**
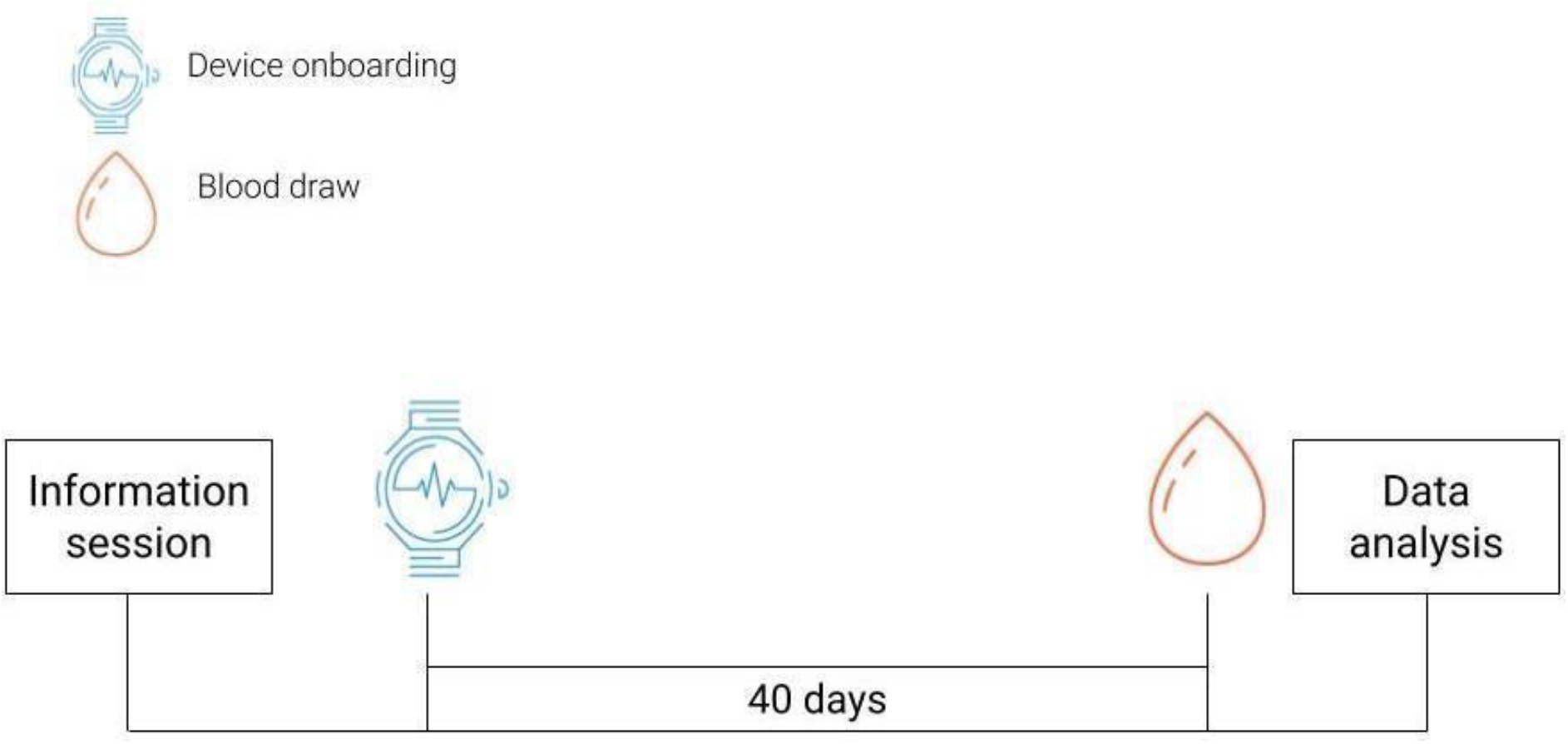
Study design

### Wearable data collection

Tri-axial accelerometer and PPG signal data were sampled at 25Hz. Thereafter signal quality filtering and feature generation occurred on both the wearable device via LifeQ’s device-enablement software libraries and then uploaded into LifeQ’s cloud environment. Study researchers were given randomized research identification codes which associated with the study ID number of the participant to map the wearable data to the blood-derived and clinical data. All data were anonymized and stored on Amazon Web Services simple storage service (AWS S3) in compliance with local data privacy legislation.

### Wearable feature analysis

Daily means were calculated for all features, after which the 40-day mean was calculated from the daily means. Wearable features were then separated into behavioural and physiological features. Behavioural features implies that an individual has the ability to alter the feature value based on their conscious decisions. Physiological features are features which give information about the physiological state of the participant and cannot be directly regulated by conscious decision making. Sleep metrics were considered to be behavioural features as an individual has the ability to alter sleep metrics, for example total sleep time is primarily determined by when one goes to and gets out of bed.

### Blood and clinical data collection

Blood was drawn by a trained phlebotomist at the Stellenbosch PathCare, a pathology laboratory. Participants were instructed to be in a fasted state and blood draws were scheduled between 08h00 – 10h00 for all participants. 20mL of blood was drawn. Multiple tubes were used depending on the analysis needed; specifically, EDTA tubes were used to collect blood needed for DNAm processing. The EDTA tube was immediately put on ice and either transported directly to the Centre of Proteomic and Genomic Research (CPGR) in Cape Town or stored in a –20° C freezer for short term storage before being transported to the CPGR. Following the blood draw, clinical measurements such as height, weight, and blood pressure were taken.

### Methylation analysis

Methylation analysis was performed by the CPGR in Cape Town. DNA from whole blood preserved in EDTA tubes was extracted using the MagMax DNA Multi-Sample Ultra 2.0 along with the KingFisher Flex Purification system. DNA quality was assessed using a Nanodrop for DNA concentrations, as well as the A 260/280 ratios. The Qubit BR dsDNA system was used for measuring dsDNA concentration. The Infinium HD workflow was used for processing the samples (https://www.illumina.com/Documents/products/workflows/), which included multiple steps to prepare the DNA through amplification, fragmentation, precipitation, resuspension, hybridisation, washing, and finally staining of the bead chip for imaging on the iScan instrument. The EPIC chip (Illumina, San Diego, CA, USA) requires an additional bisulphite conversion step prior to the Infinium workflow, which was performed using the EZ DNA Methylation Kit (Zymo Research). Detailed description of the protocol can be found in the supplementary methods. Raw .idat files were returned by the CPGR for analysis. Analysis of the methylation data was done using the R software v4.2.2 (R Foundation for Statistical Computing; https://www.r-project.org)^40^. The Chip Analysis Methylation Pipeline (ChAMP) package^41^ was used to process the beta values. Raw red and green .idat files were loaded along with wearable data for each sample as the phenotypic data and beta values were extracted and normalized using the BMIQ method^42^.

### Biological age calculation

Phenotypic age was calculated as described by Morgan Levine^43^. The PC versions of multiple DNAm ages were calculated using open-source libraries provided by Higgins-Chen and Morgan Levine^29^. Calculated DNAm ages were refined versions of Horvath^11^, Hannum^12^, GrimAge^13^ and PhenoAge^14^. Age acceleration was defined as the residual between the linear model of biological age regressed onto chronological age. Wearable features were correlated to biological age and age accelerations using the Pearson method. Significant correlations were corrected for multiple hypothesis testing using the Benjamini and Hochberg method^44^. To further investigate the relationship of associations between age acceleration and wearable features, differentially methylated positions were calculated by using the ChAMP package, using age acceleration as the independent variable. The output set of DMPs were used for further downstream analysis.

### Weighted gene co-expression network analysis

WGCNA was performed with the WGCNA package^45^ in R software with default parameters for blockwise module construction. A weighted adjacency matrix was established by calculating Pearson correlations between each CpG site pair. The co-methylation similarity matrix was raised to a power β = 12 (R^2^ of 0.75) to calculate the weighted adjacency matrix. Values were used to construct a topological overlap matrix (TOM), which provided a similarity measure. The TOM was used to calculate the corresponding dissimilarity (1-TOM). Modules were created by applying average linkage hierarchical clustering to the TOM. The dynamic tree cutting algorithm (function = cutreeDynamic, deep split = 0, minimum number of genes per module = 250, cut height = 0.15) was used to detect methylation modules. The assignment of outlying genes to modules was performed using the Partitioning Around Medoids (PAM) method. DNA methylation levels of CpG sites within a module were summarized with the module eigengene (ME) value, which is the overall methylation level of CpG sites clustering in a module. To identify relevant modules for the analysis, correlations between MEs and wearable features were calculated. Module membership (MM) was determined as the correlation of the ME and the individual CpG site. Module sites were determined by limiting the MM of individual sites to r > 0.8. To assess the underlying biology of modules, CpG sites were annotated for genes using the Genome wide annotation for Human (org.Hs.eg.db) package^46^. Gene lists were input into EnrichR library^47^ for KEGG 2021 pathway enrichment. Significant pathways were selected using an adjusted p value < 0.05. Gene lists were mapped for Entrez ID’s and input into Pathview^48^ for visual analysis of the enriched pathways.

## Results

### Study participant characteristics

A total of 48 study participants were recruited to the study. The participant chronological ages ranged from 24 – 81 years with an even distribution over and under 40 years old. The cohort was evenly distributed between males (54%) and females (46%). Participant characteristics are summarized in Table 2, detailed description of participants can be found in supplementary table 1, and blood biomarkers can be found in supplementary table 2.

**Table 2:** Study participant characteristics.

|  | Over 40 years<br>(N=22) | Under 40 years<br>(N=26) | p value |
| --- | --- | --- | --- |
| **Chronological age (years)** |  |  |  |
| Mean (SD) | 57.0 (11.0) | 31.6 (4.54) | < 0.05 |
| Median [Min, Max] | 57.0 [40.0, 81.0] | 31.5 [24.0, 39.0] |  |
| **Weight (kg)** |  |  |  |
| Mean (SD) | 82.3 (16.6) | 72.0 (15.9) | < 0.05 |
| Median [Min, Max] | 79.5 [53.0, 131] | 66.5 [51.0, 110] |  |
| **Height (m)** |  |  |  |
| Mean (SD) | 1.73 (0.0898) | 1.72 (0.0776) | 0.62 |
| Median [Min, Max] | 1.75 [1.53, 1.90] | 1.74 [1.59, 1.85] |  |
| **BMI (kg/m<sup>2</sup>)** |  |  |  |
| Mean (SD) | 27.4 (4.92) | 24.2 (4.20) | < 0.05 |
| Median [Min, Max] | 26.0 [21.0, 42.0] | 22.0 [19.0, 34.0] |  |
| **Systolic blood pressure (mmHg)** |  |  |  |
| Mean (SD) | 123 (15.5) | 109 (17.2) | < 0.05 |
| Median [Min, Max] | 122 [100, 165] | 110 [80.0, 147] |  |
| **Diastolic blood pressure (mmHg)** |  |  |  |
| Mean (SD) | 77.8 (9.60) | 72.2 (10.1) | 0.056 |
| Median [Min, Max] | 77.5 [62.0, 104] | 70.0 [56.0, 98.0] |  |

DNAm ages, in years, were calculated for study participants and are displayed in Figure 3 alongside chronological age. The means and standard deviations (SD) for all ages were calculated for the study participants shown in Figure 3: chronological age (43.73 ± 15.11), PCHorvath1 (54.3 ± 12.0), PCHannum (57.2 ± 12.7), PCGrimAge (55 ± 12.8), PCPhenoAge (43.4 ± 14.9), as well as phenotypic age (39.9 ± 15.3). The range for the calculated DNAm age accelerations in years were PCHorvath1 (19.5), PCHannum (22.2), PCGrimAge (13.5), PCPhenoAge (25.9) and Phenotypic age (12.6). Biological age calculations are summarized in supplementary table 3.

**Figure 3:**
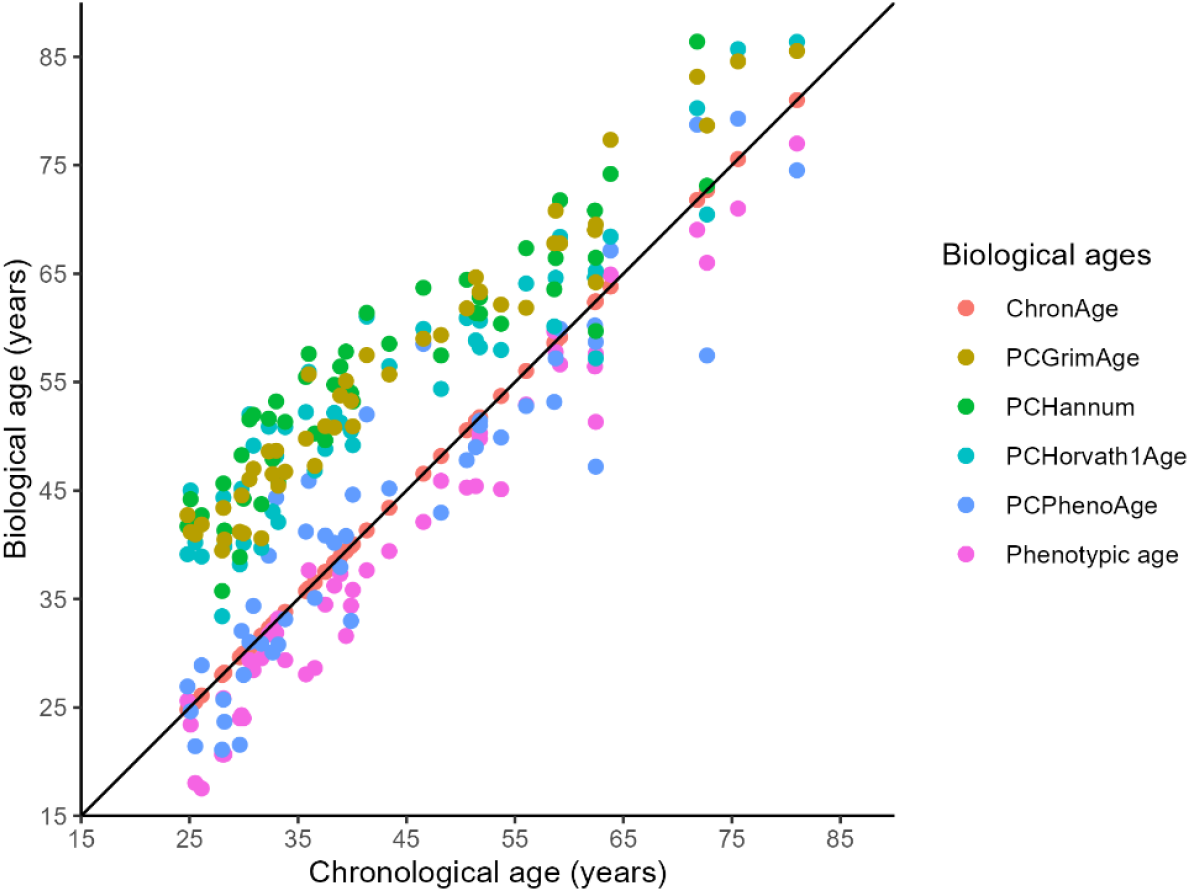
Biological age calculations. Chronological age is shown on the x-axis for each participant and the corresponding biological ages on the y-axis.

**Table 3:** Comparison of significant wearable feature and biological age correlations. Data displayed in the table are the z scores and (p values) for feature to biological age correlation.

| Feature | PCHorvath1 | PCHannum | PCPhenoAge | PCGrimAge | PhenotypicAge | ChronAge |
| --- | --- | --- | --- | --- | --- | --- |
| VO<sub>2</sub> max | -0.25 (1) | -0.28 (1) | -0.37 (0.009) | -0.25 (1) | -0.26 (1) | -0.21 (1) |
| Deep Sleep Percentage | -0.41 (1) | -0.44 (0.11) | -0.44 (0.11) | -0.38 (1) | -0.32 (1) | -0.38 (1) |
| SEVR | 0.37 (1) | 0.35 (0.63) | 0.31 (0.01) | 0.37 (1) | 0.40 (1) | 0.40 (1) |
| SAR | -0.37 (1) | -0.41 (1) | -0.47 (0.03) | -0.42 (1) | -0.44 (0.50) | -0.42 (1) |
| LASI | 0.34 (0.71) | 0.36 (1) | 0.48 (1) | 0.33 (0.71) | 0.32 (0.52) | 0.31 (0.52) |
| BPI | -0.42 (1) | -0.44 (1) | -0.46 (0.08) | -0.43 (1) | -0.46 (0.06) | -0.41 (1) |

### Wearable features

Behavioural features were correlated with physiological features (using the Pearson method); the correlations are illustrated in Figure 4, and supplementary table 4. The clustered heatmap shows two main clusters of behavioural features, clustering activity features together and sleep features with sedentary hours. Significant positive correlations between activity metrics include VO_2_ max, SEVR, and b:a ratio. Activity metrics are negatively correlated with RHR and HO. Sedentary hours are positively correlated with RHR and HO, and negatively correlated with SEVR and VO_2_ max. Deep sleep percentage is positively correlated with HO and negatively correlated with SEVR.

**Figure 4:**
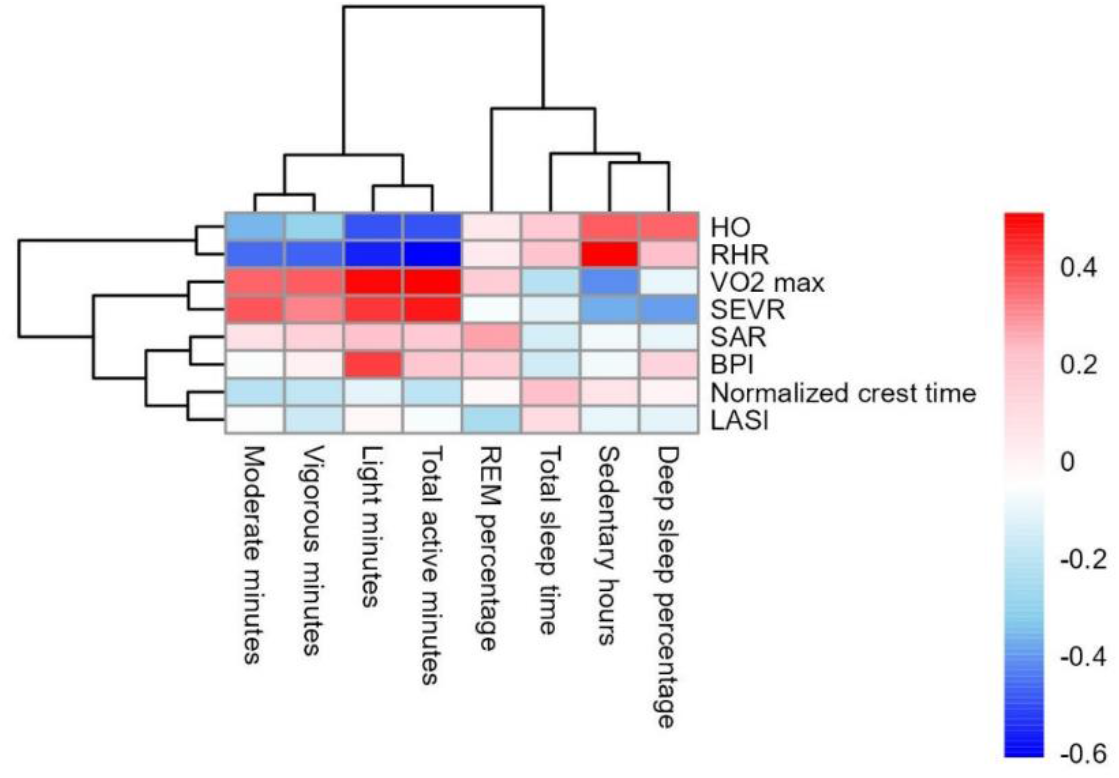
Cluster heatmap of behavioural and physiological wearable feature correlations. Behavioural and physiological features are plotted as columns and rows, respectively. The heatmap is clustered by rows and columns. Each cell represents the Pearson R value of the correlation on a sliding colour scale from negative (blue) to positive (red) correlations.

**Table 4:** Pathway enrichment for the turquoise, brown and blue modules.

| Module colour | Term | Overlap | p value | q value |
| --- | --- | --- | --- | --- |
| Turquoise | Axon guidance | 156/182 | 5.59E-07 | 1.78E-04 |
|  | Breast cancer | 126/147 | 6.81E-06 | 1.09E-03 |
|  | Oxytocin signalling pathway | 128/154 | 1.31E-04 | 1.40E-02 |
|  | Hepatocellular carcinoma | 138/168 | 2.21E-04 | 1.73E-02 |
|  | Circadian entrainment | 83/97 | 2.89E-04 | 1.73E-02 |
|  | Pathways in cancer | 407/531 | 3.26E-04 | 1.73E-02 |
|  | Human papillomavirus infection | 259/331 | 4.37E-04 | 1.99E-02 |
|  | Parathyroid hormone synthesis, secretion and action | 89/106 | 6.95E-04 | 2.77E-02 |
|  | Regulation of actin cytoskeleton | 173/218 | 1.15E-03 | 3.63E-02 |
|  | Focal adhesion | 160/201 | 1.35E-03 | 3.63E-02 |
|  | Dopaminergic synapse | 108/132 | 1.37E-03 | 3.63E-02 |
|  | Ubiquitin mediated proteolysis | 114/140 | 1.44E-03 | 3.63E-02 |
|  | Neurotrophic signalling pathway | 98/119 | 1.48E-03 | 3.63E-02 |
|  | Calcium signalling pathway | 188/240 | 2.29E-03 | 4.87E-02 |
| Brown | Human T-cell leukemia virus 1 infection | 68/219 | 3.35E-07 | 1.05E-04 |
|  | Parathyroid hormone synthesis, secretion and action | 33/106 | 3.13E-04 | 4.39E-02 |
|  | Human papillomavirus infection | 81/331 | 4.65E-04 | 4.39E-02 |
|  | Th1 and Th2 cell differentiation | 29/92 | 5.61E-04 | 4.39E-02 |
|  | Pathways in cancer | 120/531 | 7.61E-04 | 4.52E-02 |
|  | MAPK signalling pathway | 72/294 | 9.08E-04 | 4.52E-02 |
|  | FoxO signalling pathway | 37/131 | 1.13E-03 | 4.52E-02 |
|  | Axon guidance | 48/182 | 1.21E-03 | 4.52E-02 |
|  | Growth hormone synthesis, secretion and action | 34/119 | 1.41E-03 | 4.52E-02 |
|  | Notch signalling pathway | 20/59 | 1.44E-03 | 4.52E-02 |
| Blue | Human T-cell leukemia virus 1 infection | 76/219 | 5.42E-08 | 1.70E-05 |
|  | Pathways in cancer | 140/531 | 3.71E-05 | 5.82E-03 |
|  | Focal adhesion | 61/201 | 1.13E-04 | 1.18E-02 |
|  | PI3K-Akt signalling pathway | 96/354 | 2.01E-04 | 1.26E-02 |
|  | MAPK signalling pathway | 82/294 | 2.15E-04 | 1.26E-02 |
|  | Parathyroid hormone synthesis, secretion and action | 36/106 | 2.61E-04 | 1.26E-02 |
|  | Regulation of lipolysis in adipocytes | 22/55 | 3.15E-04 | 1.26E-02 |
|  | Growth hormone synthesis, secretion and action | 39/119 | 3.42E-04 | 1.26E-02 |
|  | Notch signalling pathway | 23/59 | 3.62E-04 | 1.26E-02 |
|  | Human papillomavirus infection | 89/331 | 4.55E-04 | 1.43E-02 |
|  | FoxO signalling pathway | 41/131 | 7.09E-04 | 2.02E-02 |
|  | Axon guidance | 53/182 | 9.04E-04 | 2.36E-02 |
|  | AMPK signalling pathway | 37/120 | 1.69E-03 | 3.72E-02 |
|  | Th1 and Th2 cell differentiation | 30/92 | 1.71E-03 | 3.72E-02 |
|  | Renal cell carcinoma | 24/69 | 1.81E-03 | 3.72E-02 |
|  | Thyroid hormone signalling pathway | 37/121 | 1.99E-03 | 3.72E-02 |
|  | Cholinergic synapse | 35/113 | 2.04E-03 | 3.72E-02 |
|  | Rap1 signalling pathway | 58/210 | 2.13E-03 | 3.72E-02 |
|  | Longevity regulating pathway | 32/102 | 2.47E-03 | 4.09E-02 |

### Wearable feature correlation to biological ages

Correlations were calculated between wearable features and six different age calculations: PCHorvath1, PCHannum, PCPhenoAge, PCGrimAge, Phenotypic age, and chronological age. After correcting for multiple hypothesis testing, BPI, SAR, SEVR, LASI, VO_2_ max, and deep sleep percentage were significantly correlated to one or more of the age calculations (Figure 5, supplementary table 5). To determine which feature had the strongest correlation with the age calculations, z-scores for each correlation coefficient, using the Fisher transformation, were calculated and compared (Table 3). This allowed us to identify which biological age calculation had the strongest correlation with wearable features. PCPhenoAge had the strongest correlation with VO_2_ max, SEVR, normalized pulse width, and deep sleep. Wearable features had stronger correlations to a biological age calculation, predominantly PCPhenoAge, than they did with chronological age.

**Figure 5:**
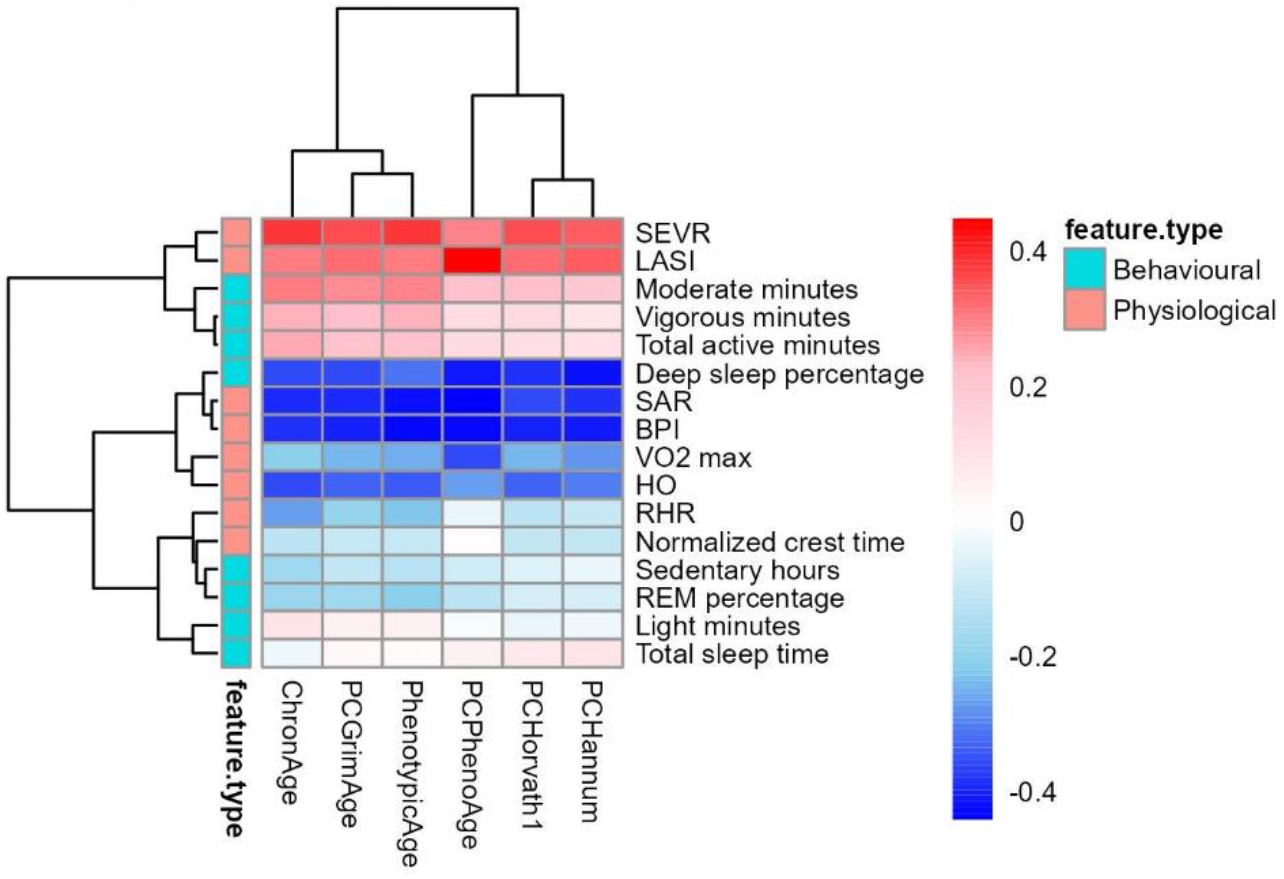
Cluster heatmap of biological age and wearable feature correlations. Biological ages are plotted as columns and wearable features as rows. Wearable features are categorized as behavioural (turquoise) and physiological (salmon). The heatmap is clustered by rows and columns. Each cell represents the Pearson R value of the correlation on a sliding colour scale from negative (blue) to positive (red) correlations.

### Wearable feature associations with age acceleration

Wearable features were correlated with biological age accelerations, displayed in the cluster heatmap in Figure 6. After adjusting the p values for multiple testing, PCPhenoAge acceleration was significantly correlated with RHR (r = 0.57, p = 1.48E-03), VO_2_ max (r = -0.45, p = 3.41E-02), and LASI (r = 0.46, p = 3.41E-02). All correlation values can be found in the supplementary table 6. The relationship between wearable features and PCPhenoAge acceleration was further explored.

**Figure 6:**
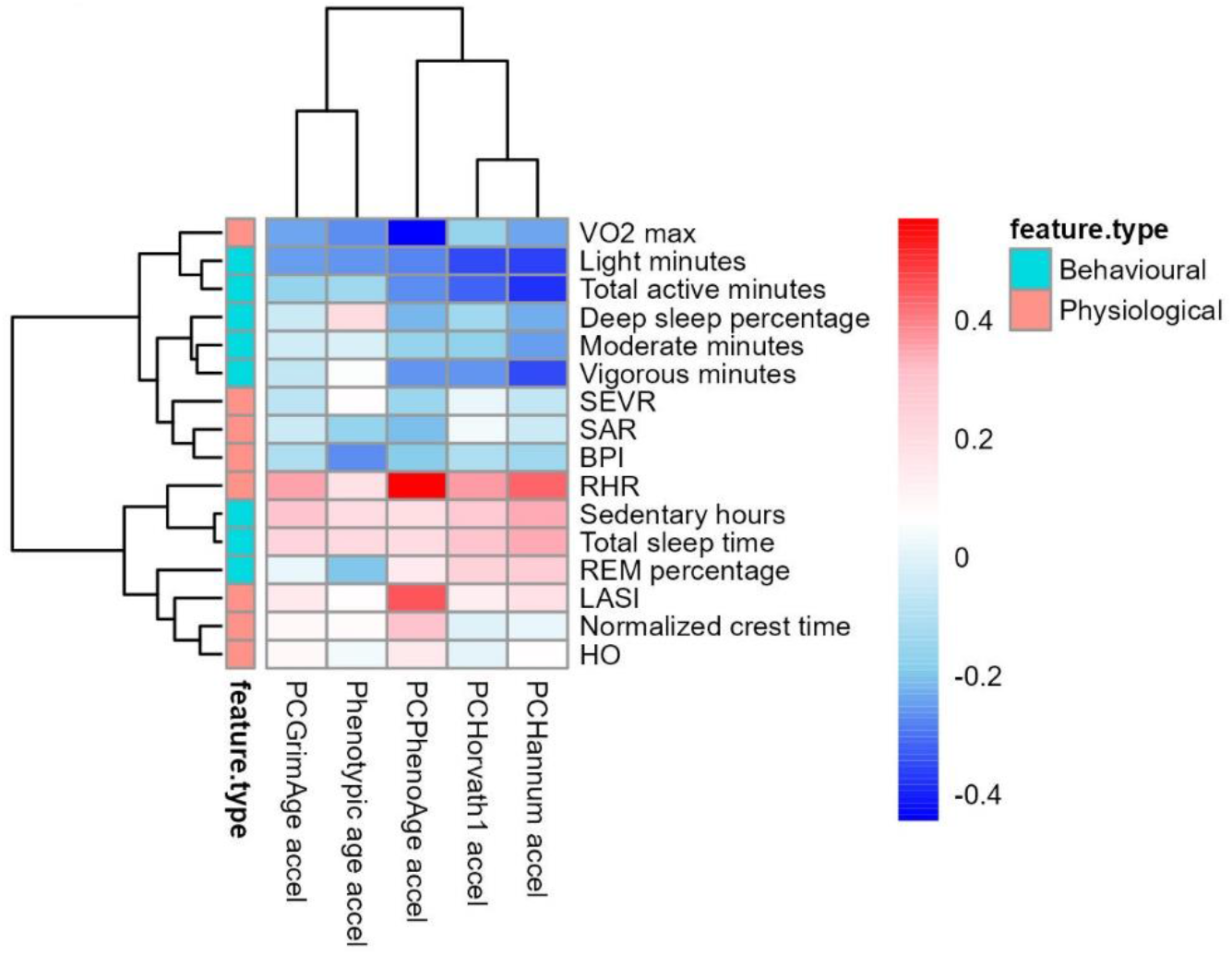
Cluster heatmap of wearable feature correlations with biological age acceleration. Biological age accelerations are plotted as columns and wearable features as rows. Wearable features are categorized as behavioural (turquoise) and physiological (salmon). The heatmap is clustered by rows and columns. Each cell represents the Pearson R value of the correlation on a sliding colour scale from negative (blue) to positive (red) correlations.

### WGCNA

Due to the significant associations discovered above, CpG site methylation levels were assessed for differentially methylated positions with PCPhenoAge acceleration as the independent variable of the linear regression using the ChAMP package. There were 65 718 DMPs as a result of the regression (adjusted p <= 0.05). This subset of CpG sites was used to construct the WGCNA modules. WGCNA produced seven modules (excluding the grey module). Module sizes are illustrated as a bar graph in Figure 7. The turquoise, blue, brown, and yellow modules were the largest modules. All modules had more than 250 CpG sites. The black module was the smallest of the modules with 251 CpG sites. The 8850 uncorrelated CpG sites formed the grey module.

**Figure 7:**
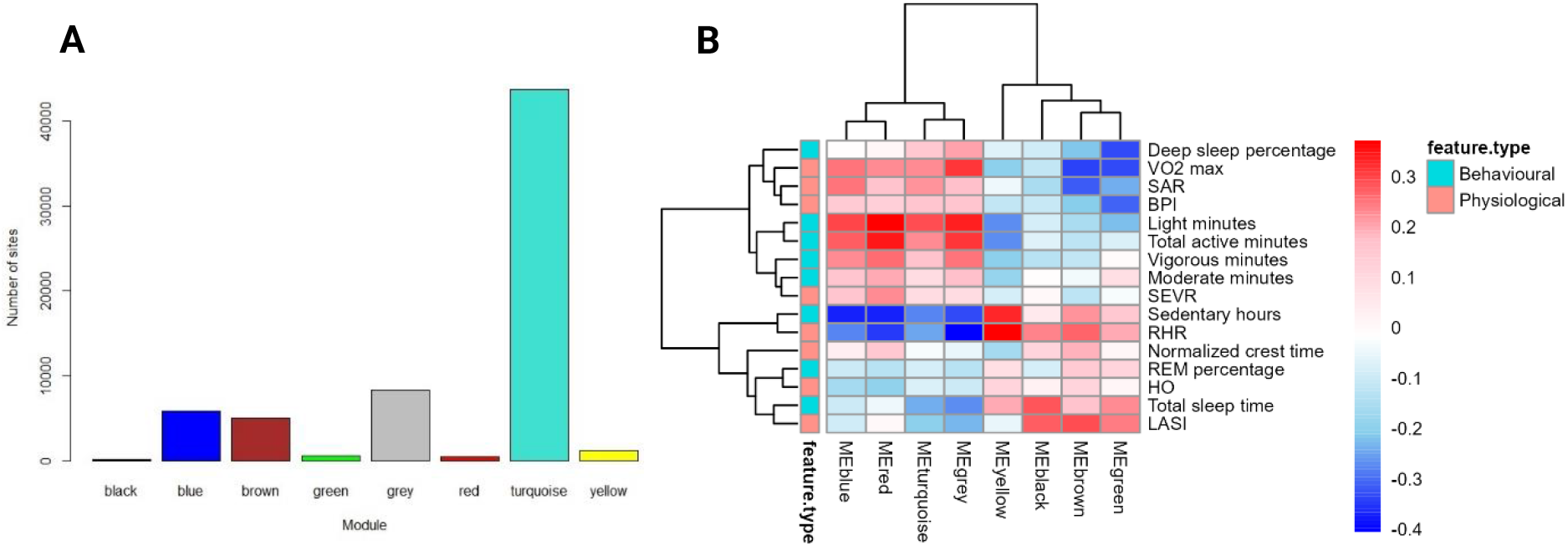
A. Bar plot showing module sizes. **B.** Cluster heatmap of features, displayed as rows, correlating with WGCNA modules eigengenes, displayed as columns. Wearable features are categorized as behavioural (turquoise) and physiological (salmon). The heatmap is clustered by rows and columns. Each cell represents the Pearson R value of the correlation on a sliding colour scale from negative (blue) to positive (red) correlations.

Wearable features (both behavioural and physiological) were correlated with the ME for each module. Before multiple hypothesis testing correction was applied, features which were significantly correlated with ME included the following: light minutes (r = 0.29, p = 0.04) and the turquoise module; light minutes (r = 0.30, p = 0.04), sedentary hours (r = -0.37, p = 0.008), RHR (r = -0.27, p = 0.05), and the blue module; the red module and sedentary hours (r = -0.37, p = 0.009), RHR (r = -0.34, p = 0.016), light minutes (r = 0.37, p = 0.009), and total active minutes (r = 0.34, p = 0.017); the yellow module and sedentary hours (r = 0.33, p = 0.021) and resting heartrate (r = 0.37, p = 0.009); and finally, the brown module and LASI (r = 0.29, p = 0.045), VO_2_ max (r = -0.34, p = 0.017), and normalized pulse width (r = -0.31, p = 0.029). After multiple testing correction these significant associations were not observed. The associations were still, however, interesting enough to further investigate the underlying biology.

### Pathway enrichment of modules of interest

The above-mentioned module CpG sites were annotated for the genes associated with the CpG sites. These gene lists formed input for pathway enrichment using the enrichR package^47^. Table 4 below illustrates the significant enrichment terms from the KEGG 2021 database. The turquoise module, being the largest, can be characterised as a module involved in regulating cell cycle, cellular homeostasis, and cellular and tissue maintenance. The brown and the blue modules seem to have a complementary relationship, as evidenced by the inverse relationship the ME had with wearable features, and the fact that both are enriched for similar pathways. Cell signalling pathways which are present in these modules are evident in many biological processes. From the enriched pathways, pathways of interest were selected for further investigation. The pathways selected for further analysis were the HTLV-1 pathway, as it is the most significantly enriched pathway in the blue and brown modules. Cell signalling pathways such as Phosphoinositide 3-kinase (PI3k) – Protein kinase B (Akt), Mitogen-activated protein kinase (MAPK), and Forkhead box O (FoxO) signalling were significantly enriched. These signalling pathways are involved in other enriched pathways, such as the longevity pathway and regulation of lipolysis in adipocytes, making them pathways of interest.

The gene list and accompanying KEGG pathway were input into pathview and resulted in KEGG pathway images (Figure 8), whereby genes coloured in red are present in the module annotated gene list. Figure 8 is an illustration of pathways and genes present in the blue module; HTLV-1 pathway of the brown module can be found in supplementary image 1. Interestingly, the CpG sites for the pathways of interest for both the blue and the brown module were annotated to the same genes. Given the complementary effect of the blue and brown modules on the same pathways, we derived a measure to illustrate the overall effect, The net ME is the arithmetic difference between the ME values of the blue and brown modules and gives a representation of the methylation status of the two modules as it pertains to the pathways of interest (Figure 9). To investigate the relationship of this differential methylation with age acceleration, the net ME values were correlated to PCPhenoAge acceleration. Although the CpG sites for the blue and brown modules were annotated to the same genes, we found that the net effect of the methylation on the pathway genes differed between individuals, and that this effect strongly correlated with PCPhenoAge acceleration (r = 0.67, p = 1.72e-07).

**Figure 8:**
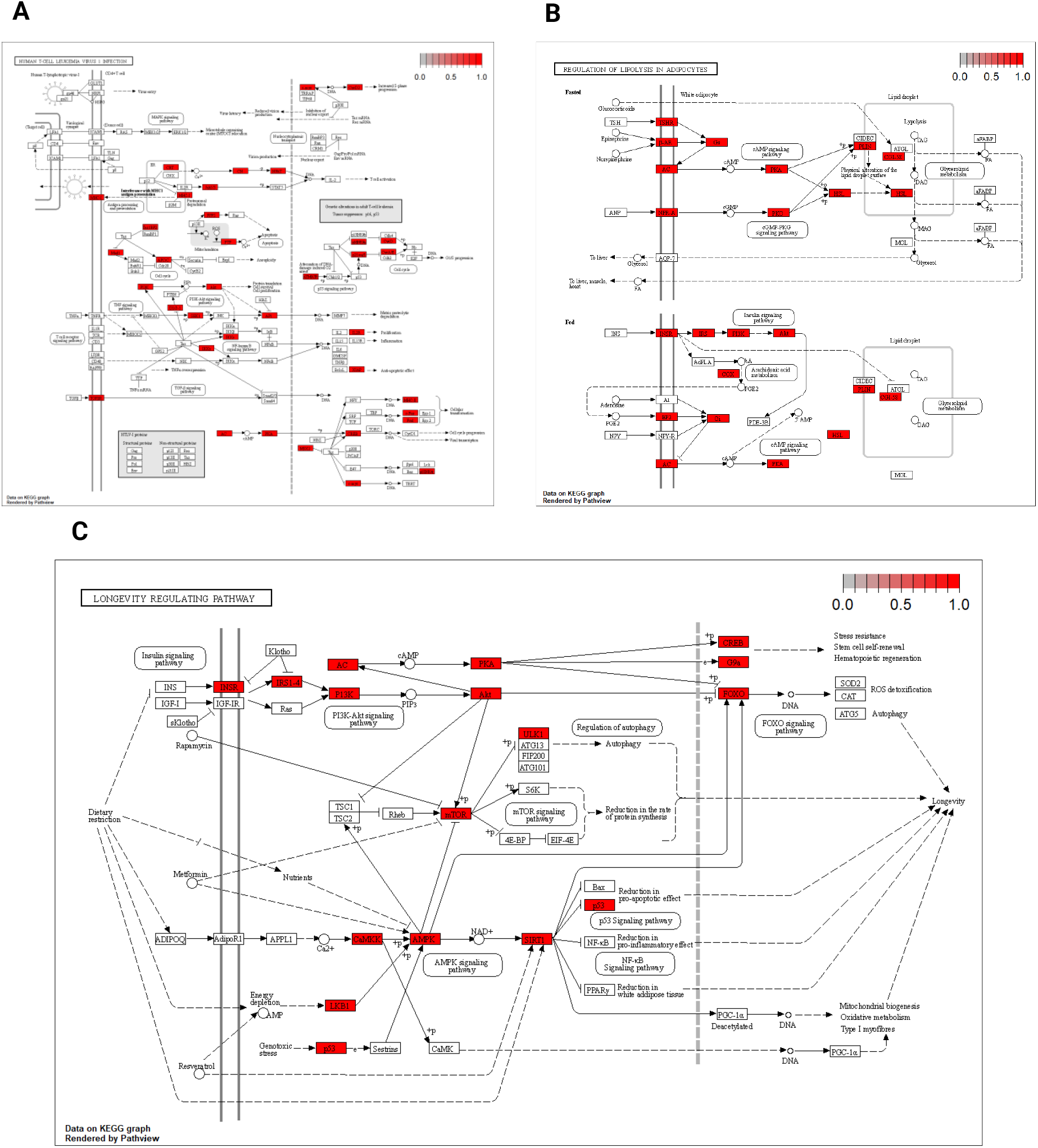
Pathways of interest annotated with genes present in the blue module. **A.** The HTLV-1 pathway. **B.** The lipolysis in adipocytes pathway. **C.** The longevity regulating pathway. Genes coloured in red indicate annotated genes found in the blue module.

**Figure 9:**
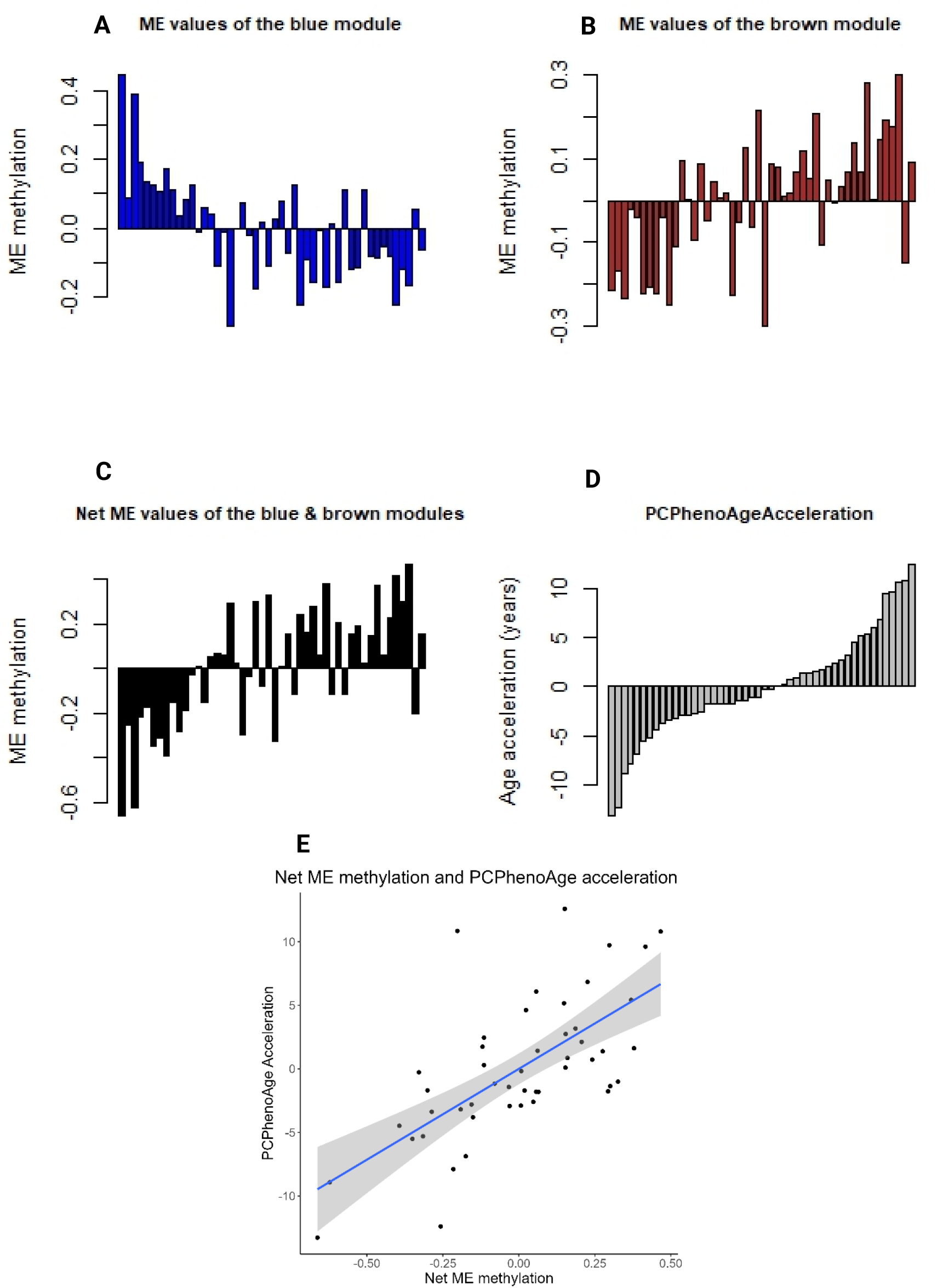
Bar plots of the individual ME values of the blue module (**A**) and brown module (**B**) ordered by ascending PCPhenoAge acceleration. **C.** Bar plot of the individual net ME values of the blue and brown modules, ordered by ascending PCPhenoAge acceleration. **D.** Bar plot of the individual PCPhenoAge accelerations. **E.** Scatter plot with regression line and confidence intervals of net ME values on the x-axis and PCPhenoAge acceleration on the y-axis.

## Discussion

Relationships between behavioural and physiological features were shown in this study. Increased activity and decreased sedentary time were related to a lower RHR and ejection duration index and higher VO_2_ max and SEVR measurements. Physiological state is dynamic, sufficient stimulus needs to be placed on the organism to elicit adaptation^49^. It is well established that lower RHR levels are associated with increased activity time^50^. The molecular mechanism driving the change in RHR is complex and multifactorial but includes parasympathetic activation, decreased responsiveness to beta-adrenergic stimulation, decreased intrinsic heart rate or a combination of all of them^51^. RHR decrease due to training has been shown to be independent of parasympathetic activation^52^. Structural changes to the heart occur including numerous growth factors such as insulin-like growth factor 1 (IGF-1), hepatocyte growth factor (HGF) and vascular endothelial growth factor (VEGF) providing a cellular environment for myocardial remodelling^53^. The exercise training is shown to increase VO_2_ max levels^54^. VO_2_ max does have a genetic component to it, but it can be altered by lifestyle behaviours. Increased activity leads to peripheral and central adaptations, such as muscle perfusion, increased mitochondrial number and function, and increased stroke volume which will increase VO_2_ max. SEVR is a measure of myocardial perfusion, blood supply to heart coronary arteries, a high SEVR indicates high blood flow to the heart. Regular physical activity can improve cardiovascular health by increasing the supply of blood to the heart muscle, as shown by a 6-week moderate intensity training program which improved SEVR in sedentary individuals^55^. The present study only captured wearable data for 40 days; this makes it difficult to characterize individual habits. A study design which includes a training intervention and longer monitoring time would aid in capturing the changes in the wearable derived features as a function of behaviour.

Considering biological age as the phenomenon of organismal function as time progresses, the stronger associations made between the wearable features and biological age, mainly PCPhenoAge, compared to that of chronological age suggests that the information captured by the wearable is closer related with the former. Features such as VO_2_ max and some of the pulse waveform features are known to be correlated with age^33,34^. and with the risk of compromised health status^56–58^. The combination of the natural change in function due to aging and the health state of the individual could explain the stronger associations of the age correlated features with biological age than chronological age. When addressing the age acceleration, this normalizes for the natural decline in function seen with age. A complete measure of age acceleration is still not agreed upon in the aging field. A technique used in this study was the use of epigenetic clocks and discovering the relationships between behavioural and physiological features and epigenetic age acceleration. One of the benefits of the use of these clocks is, by design, the output is in the scale of years. Other measures of age acceleration, such as population relativity data, are not as intuitive. Epigenetic age acceleration has been associated with an increase in ACM and disease onset^13,14,18^. What is yet to be achieved is a long-term clinical trial with multiple time points and interventions associated with decreasing aging and ACM. A study of this stature will aid in validating the use of epigenetic clocks as a measure of biological age. An increase in age acceleration, measured by the PCPhenoAge clock, was associated with high measures of RHR and LASI and a low VO_2_ max measurement. This, to our knowledge, is the first study to link epigenetic age acceleration with metrics derived from a PPG-based wearable. Another study which utilizes wearable technology to measure biological age, MoveAge, uses an accelerometer-based wearable^59^. Features derived from a combination of PPG and accelerometer data offer more biological insight to that of an accelerometer. Increased arterial stiffness, measured by impedance cardiography, has been shown to associate with an increase in epigenetic age acceleration^19^, validating our finding. Estimation of VO_2_max has been shown to negatively associate with a fitness specific DNAm clock, FitAge^60^, but not PhenoAge acceleration^61^. This speaks to a hypothesis that different epigenetic clocks capture different portions of aging. Although none of the behavioural features measured in this study associated with age acceleration, increased exercise^25,62,63^ and good sleeping habits^26,27,64^ have been shown to correlate with epigenetic age acceleration. A plausible explanation for this could be due to the small sample size (n = 48) and the duration of the study being insufficient. A study designed by Spartano *et al.* (2023), with a large sample size (n = 2435), has identified a relationship between physical activity and sedentary time, measured by an accelerometer, and epigenetic age acceleration^62^.

WGCNA is an analysis developed by Horvath and Langfelder^65^, originally for transcriptomic data, but has been used for multiple types of biological data, such as genomic^66^, proteomic^67^ and methylation data^68^. The method clusters highly connected co-methylated CpG sites into a module, represented by a ME, the first principal component of the methylation status of the sites within the module. Although following multiple hypothesis testing there were no significant associations identified between ME and wearable features, the associations found prior to the multiple hypothesis testing were worth investigating. Plausible reasons that no significant associations were found following the multiple hypothesis testing could be the small sample size, health status of the study cohort and WGCNA module generation itself. There is a large degree in variation found in human studies, this natural variation makes it difficult to find significant associations. Although the study cohort had a range of PCPhenoAge acceleration from -15 to + 16 years, a large portion of the cohort was between - 4 and + 4 years. Increasing the number of participants with age accelerations above and below 8 years may reduce the noise generated from the median age accelerated participants, as this degree of acceleration has been associated with increase in hazard ratio with ACM^14^. Finally, due to the two limitations described before this could have impacted the modules generated by WGCNA. Larger sample sizes are recommended when performing WGCNA, and adding in participants who were age accelerated and decelerated would also improve the network construction, potentially leading to significant associations with wearable features. The modules of interest were identified from the correlations between features and ME. Light minutes negatively correlated with the turquoise ME. The turquoise module, being the largest, is difficult to summarize but can be classified as a module responsible for general cellular, and tissue maintenance module. The molecular footprint of aging includes general dysregulation of cellular and tissue homeostasis and maintenance pathways^69^. A hallmark of aging is genetic instability^5^. This will increase the chance of misfolding of proteins, for example. In a healthy cell, these proteins would be recycled by the ubiquitin-mediated proteolysis and autophagy processes^70^. If these processes are attenuated or the accumulation of proteins is above the capacity of autophagic flux these accumulated proteins can be detrimental, as seen progression of Alzheimer’s disease^71^. Pathways for cancer were enriched in the turquoise module. The risk for cancer increases with age^2^ and epigenetic age acceleration^22^. Molecular interactions which lead to cancers are complex and multi-factorial. Mutations to a key cancer gene, p53, have been noted in cancer patients^72^. There is also an indirect impact on the cancer pathology whereby epigenetic modifications allow cancer cells to escape immune cell surveillance^73^. There is evidence that increased physical activity reduces the risk of cancer and that up to 40% of the risk of cancer can be mitigated through lifestyle behaviours^74^. This relationship can be inferred from this as study as moderate minutes associated with the ME of the turquoise module, validating the idea of using activity as a tool to prolong cellular integrity.

Although, the CpG sites for the blue and brown modules were annotated to the same genes, we found that the net effect of the methylation on the pathway genes differed between individuals, and that this effect strongly correlated with PCPhenoAge acceleration, allowing us to conclude that the specific methylation pattern in age-related pathways results in different aging phenotypes.

Antiviral defence mechanisms have co-evolved with humans and include methylation of CpG sites to inactivate retrovirus expression. Consequences of HTLV-1 infection are wide-ranging, with alterations in cell-cycle progression, cell proliferation and inflammation. HTLV-1 infection causes epigenetic reprogramming^75^, resulting in HTLV-1-associated myelopathy/tropical spastic paraparesis (HAM/TSP) and adult T-cell leukemia–lymphoma (ATL) as downstream events, instead of the normal CD4+ T-lymphocyte phenotype. Specifically in the case of ATL/ATLL the host genome shows high levels of hypermethylation in CpG islands in promotor regions. It is therefore notable, that virtually all genes involved in this pathway are differentially methylated with regards to PCPhenoAge acceleration, and furthermore, that the precise methylation state (hyper-/hypomethylation) is seen in two WGCNA modules that differs between individuals and correlates with individual PCPhenoAge acceleration. During HTLV-1 infection, the viral proteins Tax and HBZ are associated with wide-ranging methylation of host cell DNA^75^, altering the cell fate landscape, and ultimately leading to ATL. Having observed epigenetic changes in the same pathway in individuals without HTLV-1 infection suggests that other drivers (in this case likely human behaviour) converge on the same pathway to similarly result in potentially adverse outcomes.

Metabolic risk factors such as elevated glucose, hypertension, increased BMI and insulin resistance are associated with epigenetic age acceleration^17^. These risk factors make up the metabolic syndrome^76^. Lipolysis in adipocytes is the process which hydrolyses triglycerides (TAG) to release free fatty acids (FFA) and glycerol into circulation^77^. The regulation of this process has 3 main signalling pathways, cAMP, insulin and guanosine 3’,5’-cyclic monophosphate (cGMP) signalling pathways. Upon binding of glucagon or epinephrine, cyclic adenosine monophosphate (cAMP) and consequently protein kinase A (PKA) are activated. PKA activates hormone sensitive lipase (HSL) which is responsible for breaking down TAG to FFA. Protein kinase G (PKG) acts on HSL in a similar fashion to PKA in the cGMP pathway. The insulin signalling pathway involves binding of insulin and a cellular cascade which activates PI3K/Akt which inhibits the translocation of HSL from the cytoplasm to the surface of the lipid droplet, inhibiting lipolysis. Additionally, insulin increases the uptake of glucose in adipocytes for storage. CpG sites annotated to genes in all three of these pathways were differentially methylated in this study. Dysregulation of lipolysis in adipocytes is seen in the aging phenotype, leading to lipid synthesis and ultimately increased adipose tissue as a structural change, but indirectly increases chronic inflammation accompany this dysregulation. Increased physical activity improves insulin sensitivity^78^ and may attenuate the effects of dysfunctional lipolysis in adipocytes.

Genes involved in the previously mentioned pathway as INSR/IRS/PI3k/Akt occur in multiple cellular processes such as cellular metabolism and cell survival^79^, but of interest in this study is their enrichment in the longevity regulating pathway. Maintaining cellular homeostasis is paramount to cellular health and function. Multiple signalling pathways are responsible for this such as PI3k/Akt/FoxO, PI3k/Akt/mTOR (mechanistic target of rapamycin) and the Sirtuins. PI3k/Akt/mTOR is a signalling pathway critical in regulating cellular growth and survival^79^. mTOR can be activated by insulin and other growth signals. Under anabolic conditions, mTOR is responsible for protein and lipid synthesis, and the inhibition of autophagy. In low nutrient environments, mTOR is inhibited and autophagy is upregulated. Dysregulation of this process leads to an accumulation of damaged cells, a hallmark of aging^5^. In mice models, inhibition of this pathway has shown to improve age-related phenotypes and increased lifespan^80^. The Sirtuin family includes seven proteins which localize at various cellular compartments and regulates epigenetic modifications and cellular metabolism^81^. Differentially methylated genes in this study highlighted the interaction between 5’ AMP-activated protein kinase (AMPK) and SIRT1. AMPK is a cellular energy sensor that is activated in response to low energy states, while SIRT1 is a NAD+ dependent deacetylase. AMPK and SIRT1 interact with one another and are two role players in the aging process^82^. AMPK activation stimulates SIRT1 activity by increasing the cellular NAD+/NADH ratio, required for SIRT1 deacetylase activity^83^. A reduction in SIRT1 activation has been linked to attenuation of mitochondrial biogenesis, DNA repair and cellular stress resistance^82^. These Sirtuins have furthermore been proposed in the loss of information theory of aging. This theory suggests that an accumulation of errors in genetic information occurs over time, ultimately leading to cellular and tissue dysfunction^84^. SIRT1 has been shown to regulate base excision repair and homologous recombination^85^. Sirtuins also regulate other cellular processes and it is theorised that these molecules regulate those processes and cannot maintain genomic stability, leading to progression of the aging phenotype^82^. Interventions in mice, both pharmaceutical^86–88^ and behavioural^89^, have been shown to upregulate sirtuin activity allowing the molecules to return to maintaining genomic stability, ultimately reversing the effects of aging. SIRT3 levels have been shown to be elevated in human muscle biopsy samples in endurance trained individuals, regardless of age, compared with sedentary individuals, suggesting that exercise may attenuate an age-related decline in SIRT3 levels^90^. The associations in this study with sedentary hours and the blue and brown MEs suggest a connection between activity and methylation of the SIRT1 gene. Inactive mice have decreased sirtuin activity and present with an aging phenotype. LASI, normalized pulse width, RHR, VO_2_ max associated with the ME of the blue and brown modules. This association validates a connection between differential methylation of genes involved in molecular mechanisms of aging and physiological and structural state of humans.

Our analysis highlights a new hypothesis, which is that by overcoming the study limitations listed below, it may be possible to associate changes in wearable derived behaviour to changes in the activity of individual cellular components, especially through continued epigenomic sampling combined with the use of a wearable device. For components with a known cellular function, such a finding can potentially advance cause-and-effect understanding of the impact of lifestyle on age acceleration, going beyond purely associative relationships. We note that investigation of the transcriptome to confirm predicted effects of methylation changes might in future help further extended findings towards causal mechanisms.

### Study limitations and future recommendations

Human research is accompanied by natural variation amongst the participants of the study cohort. Increasing the sample size aids in limiting the variation observed. A limitation to this study was the small sample size, when comparing analysis techniques used by other studies. The study design was cross-sectional, only allowing for associations to be made between metrics measured. A longitudinal study with multiple measures of epigenetic age will provide an opportunity for assessing the changes in biological age over time and how the wearable is able to track those changes. Additionally, it will be interesting to assess to what degree wearable features respond to interventions which are known to slow down the rate of aging. This study did not target the recruitment of individuals with compromised health. Adding these individuals to future studies would make the observed associations more robust and allow the analysis to group the participants, allowing causal relationships to be discovered.

## Conclusion

Behavioural features impact physiological state, measured by the wearable. Wearable derived physiological features give an indication of health that is more strongly correlated with biological age than chronological age and correlated with age acceleration. These physiological features are a by-product of behavioural features, for example, increased activity time and decreased sedentary time are linked to improved physiological metrics. Exploration of the underlying biology of age acceleration presented trend correlations with behavioural and physiological features and methylation of genes in pathways relating to the aging process. This study presents the case that PPG-based wearable devices can capture portions of biological aging and should be used as a tool, in combination with other measurement techniques, for monitoring and potentially, in future, helping to manage healthy aging.

## Supporting information

supplementary image 1

## Acknowledgements

Members of the LifeQ team assisted in the data collection process for this study. Special mention must be made to Lara Grobler, Sarah Arnold, Maia Rawlins, Shona Troost, Wian Smit, and Corné Coetzee in their roles assisting with the data collection. Ongoing intellectual support as this study progressed was exhibited by Alexa Lewis and Jianhua Ma. Nurses at PathCare Stellenbosch and scientists from the CPGR are acknowledged for the drawing and processing the blood samples respectively.

## References

1. Aunan, J. R., Watson, M. M., Hagland, H. R. & Søreide, K. Molecular and biological hallmarks of ageing. British Journal of Surgery 103, e29–e46 (2016).

2. Kaeberlein, M. Longevity and aging. F1000Prime Rep 5, 1–8 (2013).

3. Petsko, G. A. A seat at the table. Genome Biol 9, 113 (2008).

4. Seals, D. R., Justice, J. N. & Larocca, T. J. Physiological geroscience: Targeting function to increase healthspan and achieve optimal longevity. Journal of Physiology 594, 2001–2024 (2016).

5. López-Otín, C., Blasco, M. A., Partridge, L., Serrano, M. & Kroemer, G. The Hallmarks of Aging Europe PMC Funders Group. Cell 153, 1194–1217 (2013).

6. Komen, J. C. & Thorburn, D. R. Modeling mitochondrial dysfunction in neurodegenerative disease. Mitochondrial Dysfunction in Neurodegenerative Disorders 1366, 193–212 (2012).

7. Stojanovic, S. D., Fiedler, J., Bauersachs, J., Thum, T. & Sedding, D. G. Senescence-induced inflammation: An important player and key therapeutic target in atherosclerosis. Eur Heart J 41, 2983–2996 (2020).

8. Jones, M. J., Goodman, S. J. & Kobor, M. S. DNA methylation and healthy human aging. Aging Cell 14, 924–932 (2015).

9. Seale, K., Horvath, S., Teschendorff, A., Eynon, N. & Voisin, S. Making sense of the ageing methylome. Nat Rev Genet 23, 585–605 (2022).

10. Horvath, S. & Raj, K. DNA methylation-based biomarkers and the epigenetic clock theory of ageing. Nat Rev Genet 19, 371–384 (2018).

11. Horvath, S. DNA methylation age of human tissues and cell types. Genome Biol 14, (2013).

12. Hannum, G. et al. Genome-wide Methylation Profiles Reveal Quantitative Views of Human Aging Rates. Mol Cell 49, 359–367 (2013).

13. Lu, A. T. et al. DNA methylation GrimAge strongly predicts lifespan and healthspan. Aging 11, 303–327 (2019).

14. Levine, M. E. et al. An epigenetic biomarker of aging for lifespan and healthspan. bioRxiv 10, 573–591 (2018).

15. Liu, Z. et al. Erratum: A new aging measure captures morbidity and mortality risk across diverse subpopulations from NHANES IV: A cohort study (PLoS Med (2018) 15:12 (e1002718) DOI: 10.1371/journal.pmed.1002718). PLoS Med 16, 1–20 (2019).

16. Rezwan, F. I. et al. Association of adult lung function with accelerated biological aging. Aging 12, 518–542 (2020).

17. Ammous, F. et al. Epigenetic age acceleration is associated with cardiometabolic risk factors and clinical cardiovascular disease risk scores in African Americans. Clin Epigenetics 13, 1–13 (2021).

18. Perna, L. et al. Epigenetic age acceleration predicts cancer, cardiovascular, and all-cause mortality in a German case cohort. Clin Epigenetics 8, 1–7 (2016).

19. Liu, D., Aziz, N. A., Pehlivan, G. & Breteler, M. M. B. Cardiovascular correlates of epigenetic aging across the adult lifespan: a population-based study. Geroscience (2023) doi:10.1007/s11357-022-00714-0.

20. Ritz, B. R. & Horvath, S. Increased epigenetic age and granulocyte counts in the blood of Parkinson’s disease patients. Aging 7, 1130–1142 (2015).

21. Pellegrini, C. et al. A Meta-Analysis of Brain DNA Methylation Across Sex, Age, and Alzheimer’s Disease Points for Accelerated Epigenetic Aging in Neurodegeneration. Front Aging Neurosci 13, 1–21 (2021).

22. Durso, D. F. et al. Acceleration of leukocytes’ epigenetic age as an early tumorand sex-specific marker of breast and colorectal cancer. Oncotarget 8, 23237–23245 (2017).

23. Okazaki, S. et al. Decelerated epigenetic aging associated with mood stabilizers in the blood of patients with bipolar disorder. Transl Psychiatry 10, (2020).

24. Katrinli, S. et al. Evaluating the impact of trauma and PTSD on epigenetic prediction of lifespan and neural integrity. Neuropsychopharmacology 45, 1609–1616 (2020).

25. Quach, A. et al. Epigenetic clock analysis of diet, exercise, education, and lifestyle factors. Agro Food Ind Hi Tech 29, 20–21 (2018).

26. Li, X. et al. Association between sleep disordered breathing and epigenetic age acceleration: Evidence from the Multi-Ethnic Study of Atherosclerosis. EBioMedicine 50, 387–394 (2019).

27. Carskadon, M. A. et al. A pilot prospective study of sleep patterns and DNA methylation-characterized epigenetic aging in young adults. BMC Res Notes 12, 583 (2019).

28. Sugden, K. et al. Patterns of Reliability: Assessing the Reproducibility and Integrity of DNA Methylation Measurement. Patterns 1, 100014 (2020).

29. Higgins-chen, A. T. et al. clocks: Implications for clinical trials and longitudinal tracking. 2, 644–661 (2022).

30. Griffin, P. Ultra-cheap and scalable epigenetic age predictions with TIME-Seq. (2021) doi:10.1101/2021.10.25.465725.

31. Iconaru, E. I., Ciucurel, M. M., Georgescu, L. & Ciucurel, C. Hand grip strength as a physical biomarker of aging from the perspective of a Fibonacci mathematical modeling. BMC Geriatr 18, 1–9 (2018).

32. Habibi, E., Dehghan, H., Moghiseh, M. & Hasanzadeh, A. Study of the relationship between the aerobic capacity (VO2 max) and the rating of perceived exertion based on the measurement of heart beat in the metal industries Esfahan. J Educ Health Promot 3, 55 (2014).

33. Betik, A. C. & Hepple, R. T. Determinants of VO2 max decline with aging: An integrated perspective. Applied Physiology, Nutrition and Metabolism 33, 130–140 (2008).

34. McEniery, C. M. et al. Normal vascular aging: Differential effects on wave reflection and aortic pulse wave velocity - The Anglo-Cardiff Collaborative Trial (ACCT). J Am Coll Cardiol 46, 1753– 1760 (2005).

35. Yang, J. et al. Analysis of the Radial Pulse Wave and its Clinical Applications: A Survey. IEEE Access 9, 157940–157959 (2021).

36. Gartstein. Photoplethysmography Sensors and Their Potential Future Applications in Health Care. Physiol Behav 176, 139–148 (2016).

37. Allen, J. Photoplethysmography and its application in clinical physiological measurement. Physiol Meas 28, (2007).

38. Matthews, C. E., Hagströmer, M., Pober, D. M. & Bowles, H. R. Best Practices for Using Physical Activity Monitors. Med Sci Sports Exerc. 44, 1–17 (2012).

39. Secrest, A. M., Marshall, S. L., Miller, R. G., Prince, C. T. & Orchard, T. J. Pulse wave analysis and cardiac autonomic neuropathy in type 1 diabetes: A report from the pittsburgh epidemiology of diabetes complications study. Diabetes Technol Ther 13, 1264–1268 (2011).

40. R Core. R: A language and environment for statistical computing. R Foundation for Statistical Computing, Vienna, Austria. URL https://www.R-project.org/. Preprint at (2021).

41. Tian, Y., Morris, T., Stirling, L., Feber, A. & Teschendorff, A. Chip Analysis Methylation Pipeline for Illumina HumanMethylation450 and EPIC. Preprint at (2019).

42. Teschendorff, A. E. et al. A beta-mixture quantile normalization method for correcting probe design bias in Illumina Infinium 450 k DNA methylation data. Bioinformatics 29, 189–196 (2013).

43. Liu, Z. et al. Phenotypic Age: A novel signature of mortality and morbidity risk. bioRxiv (2018) doi:10.1101/363291.

44. Benjamini, Y. Discovering the false discovery rate. J R Stat Soc Series B Stat Methodol 72, 405– 416 (2010).

45. Zhang, B. & Horvath, S. A general framework for weighted gene co-expression network analysis. Stat Appl Genet Mol Biol 4, (2005).

46. Carlson, M. org.Hs.eg.db: Genome wide annotation for Human. Preprint at (2022).

47. Jawaid, W. enrichR. Preprint at https://maayanlab.cloud/Enrichr/ (2022).

48. Luo, W. Pathview: a tool set for pathway based data integration and visualization. Preprint at https://github.com/datapplab/pathview (2023).

49. Hedayatpour, N. & Falla, D. Physiological and Neural Adaptations to Eccentric Exercise: Mechanisms and Considerations for Training. Biomed Res Int 2015, (2015).

50. Reimers, A. K., Knapp, G. & Reimers, C. D. Effects of exercise on the resting heart rate: A systematic review and meta-analysis of interventional studies. J Clin Med 7, (2018).

51. Matsui, K. & Sugano, S. Relation of intrinsic heart rate and autonomic nervous tone to resting heart rate in the young and the adult of various domestic animals. Nippon juigaku zasshi. The Japanese journal of veterinary science 51, 29–34 (1989).

52. Stratton, J. R. Exercise Training Bradycardia Is Largely Explained. Int J Cardiol 213–216 (2017) doi:10.1016/j.ijcard.2016.07.203.EXERCISE.

53. Qiu, Y., Pan, X., Chen, Y. & Xiao, J. Hallmarks of exercised heart. J Mol Cell Cardiol 164, 126– 135 (2022).

54. Tangen, E. M., Gjestvang, C., Stensrud, T. & Haakstad, L. A. H. Is there an association between total physical activity level and VO2max among fitness club members? A cross-sectional study. BMC Sports Sci Med Rehabil 14, 1–8 (2022).

55. Tang, S. et al. Effects of aquatic high-intensity interval training and moderate-intensity continuous training on central hemodynamic parameters, endothelial function and aerobic fitness in inactive adults. J Exerc Sci Fit 20, 256–262 (2022).

56. Lee, C. Do, Blair, S. N. & Jackson, A. S. Cardiorespiratory fitness, body composition, and all-cause and cardiovascular disease mortality in men. American Journal of Clinical Nutrition 69, 373–380 (1999).

57. Sui, X., Sarzynski, M. A., Gribben, N., Zhang, J. & Lavie, C. J. Cardiorespiratory Fitness and the Risk of All-Cause, Cardiovascular and Cancer Mortality in Men with Hypercholesterolemia. J Clin Med 11, (2022).

58. Abdullah Said, M., Eppinga, R. N., Lipsic, E., Verweij, N. & van der Harst, P. Relationship of arterial stiffness index and pulse pressure with cardiovascular disease and mortality. J Am Heart Assoc 7, (2018).

59. McIntyre, R. L., Rahman, M., Vanapalli, S. A., Houtkooper, R. H. & Janssens, G. E. Biological Age Prediction From Wearable Device Movement Data Identifies Nutritional and Pharmacological Interventions for Healthy Aging. Frontiers in Aging 2, 1–11 (2021).

60. McGreevy, K. M. et al. DNAmFitAge: biological age indicator incorporating physical fitness. Aging 15, 1–35 (2023).

61. Matyas Jokai, Ferenc Torma, Kristen M. McGreevy, Erika Koltai, Zoltan Bori, Gergely Babszki, Peter Bakonyi, Zoltan Gombos, Bernadett Gyorgy, Dora Aczel, Laszlo Toth, Peter Osvath, Marcell Fridvalszky, Timea Teglas, Balazs Ligeti, Regina Kalcsevszki, Aniko, Z. R. DNA methylation clock DNAmFitAge shows regular exercise is associated with slower aging and systemic adaptation. medRxiv 27, 58–66 (2022).

62. Spartano, N. L. et al. Association of Accelerometer-Measured Physical Activity and Sedentary Time with Epigenetic Markers of Aging. Med Sci Sports Exerc 55, 264–272 (2023).

63. Kresovich, J. K. et al. Associations of Body Composition and Physical Activity Level With Multiple Measures of Epigenetic Age Acceleration. Am J Epidemiol 190, 984–993 (2021).

64. Carroll, J. E. et al. Postpartum sleep loss and accelerated epigenetic aging. Sleep Health 7, 362–367 (2021).

65. Langfelder, P. & Horvath, S. WGCNA: An R package for weighted correlation network analysis. BMC Bioinformatics 9, (2008).

66. Tian, Z. et al. Identification of important modules and biomarkers in breast cancer based on WGCNA. Onco Targets Ther 13, 6805–6817 (2020).

67. Pei, G., Chen, L. & Zhang, W. WGCNA Application to Proteomic and Metabolomic Data Analysis. Methods in Enzymology vol. 585 (Elsevier Inc., 2017).

68. Tremblay, B. L., Guénard, F., Lamarche, B., Pérusse, L. & Vohl, M. C. Network analysis of the potential role of DNA methylation in the relationship between plasma carotenoids and lipid profile. Nutrients 11, (2019).

69. Hartl, F. U. Cellular Homeostasis and Aging. Annu Rev Biochem 85, 1–4 (2016).

70. Pohl, C. & Dikic, I. Cellular quality control by the ubiquitin-proteasome system and autophagy. Science (1979) 366, 818–822 (2019).

71. Watanabe, Y., Taguchi, K. & Tanaka, M. Ubiquitin, Autophagy and Neurodegenerative Diseases. Cells 9, 1–15 (2020).

72. Hu, J. et al. Targeting mutant p53 for cancer therapy: direct and indirect strategies. J Hematol Oncol 14, 1–19 (2021).

73. Cao, J. & Yan, Q. Cancer Epigenetics, Tumor Immunity, and Immunotherapy. Trends Cancer 6, 580–592 (2020).

74. Friedenreich, C. M., Ryder-Burbidge, C. & McNeil, J. Physical activity, obesity and sedentary behavior in cancer etiology: epidemiologic evidence and biologic mechanisms. Mol Oncol 15, 790–800 (2021).

75. Yamagishi, M., Fujikawa, D., Watanabe, T. & Uchimaru, K. HTLV-1-mediated epigenetic pathway to adult T-cell leukemia-lymphoma. Front Microbiol 9, 1–8 (2018).

76. Huang, P. L. A comprehensive definition for metabolic syndrome. DMM Disease Models and Mechanisms 2, 231–237 (2009).

77. Ahmadian, Maryam (Departement of Nutritional Science and Toxicology, University of California, Berkeley, California 94720) Wang, Yuhui (Departement of Nutritional Science and Toxicology, University of California, Berkeley, California 94720), Sul, Hei Sook, C. 94720). Medicine in Focus: Lipolysis in Adipocytes. Int J Biochem Cell Biol 42, 555–559 (2010).

78. Lin, Y. et al. The Association Between Physical Activity and Insulin Level Under Different Levels of Lipid Indices and Serum Uric Acid. Front Physiol 13, (2022).

79. Hemmings, B. A. & Restuccia, D. F. PI3K-PKB / Akt Pathway. Cold Spring Harb Perspect Med 4, 1–4 (2012).

80. Harrison, D. E., et al. Heterogeneous Mice. Nature 460, 392–395 (2009).

81. Nakagawa, T. & Guarente, L. Sirtuins at a glance. J Cell Sci 124, 833–838 (2011).

82. Imai, S. ichiro & Guarente, L. NAD+ and sirtuins in aging and disease. Trends Cell Biol 24, 464– 471 (2014).

83. Cantó, C. et al. AMPK regulates energy expenditure by modulating NAD + metabolism and SIRT1 activity. Nature 458, 1056–1060 (2013).

84. Yang, J. H. et al. Loss of epigenetic information as a cause of mammalian aging. Cell 186, 305–326.e27 (2023).

85. Roos, W. P. & Krumm, A. The multifaceted influence of histone deacetylases on DNA damage signalling and DNA repair. Nucleic Acids Res 44, 10017–10030 (2016).

86. Gu, X. S. et al. Resveratrol, an activator of SIRT1, upregulates AMPK and improves cardiac function in heart failure. Genetics and Molecular Research 13, 323–335 (2014).

87. Qin, X. et al. Metformin prevents murine ovarian aging. Aging 11, 3785–3794 (2019).

88. Lázaro, I., Cossu, G. & Kostarelos, K. Transient transcription factor (OSKM) expression is key towards clinical translation of in vivo cell reprogramming. EMBO Mol Med 9, 733–736 (2017).

89. Hokari, F. et al. Muscle contractile activity regulates Sirt3 protein expression in rat skeletal Muscles. J Appl Physiol 109, 332–340 (2010).

90. Lanza, I. R. et al. Endurance exercise as a countermeasure for aging. Diabetes 57, 2933–2942 (2008).

