## supplementary methods for "Wearable-ome meets epigenome: A novel approach to measuring biological age with wearable devices"

The project will be run in accordance with this ASP and the standard terms and conditions of the CPGR. Either the Qiagen Mini DNA extraction kit or the MagMax DNA Multi-Sample Ultra 2.0 Kit along with the KingFisher™ Flex Purification system will be used to extract genomic DNA from the whole blood samples in accordance with the manufacturer's instructions. After DNA extraction, the DNA will be quantified to determine the concentration, quality, and integrity to ensure the below criteria are met for the stipulated assay:

1. Nanodrop readings for DNA concentration, as well as the $A_{260/280}$ and $A_{260/230}$ ratios. The $A_{260/280}$ ratio should be between 1.8 and 2.1 and the $A_{260/230}$ ratio should be be tween 1.5 and 3.0.
2. Either the QuantiFluor dsDNA quantification assay or the Qubit BR dsDNA system may be used for dsDNA concentration determination. It would be ideal if the dsDNA concentration exceeds 50 ng/ul, however, as long as a yield of 1000 ng of DNA re mains for processing after QC analysis has been completed, the sample will be deemed to have a high sufficient yield.
3. A 2% Agarose gel with a suitable ladder to confirm High Molecular Weight DNA.

The samples must have high molecular weight DNA, good purity ratios with a minimum yield of 1000 ng to proceed with the array processing. If samples do not meet the minimum required quality control (QC) standards specified in the ASP, the CPGR can proceed with the analysis and process such samples, but a disclaimer letter will be required as authorization. Should any of these samples not perform optimally on the array used for the project, the CPGR will not be held responsible for the outcome.

The Infinium HD workflow will be used for processing of the samples, (https://www.illumina.com/Documents/products/workflows/ workflow_infinium_ii.pdf) which includes multiple steps to prepare the DNA through amplification, fragmentation, precipitation, resuspension, hybridisation, washing, and finally staining of bead chip for imaging on the iScan instrument. The EPIC methylation array requires an additional bisulphite conversion step prior to the Infinium workflow, which will be performed using the EZ DNA Methylation Kit by Zymo Research. Any deviations will be noted and reported in a timely fashion. The CPGR will use the Methylation Module of GenomeStudio Software 2011.1 and/or the BeadArray Controls Reporter Installer Software for assessing the quality of the data produced. The Methylation Module of GenomeStudio Software 2011.1 and the BeadArray Controls Reporter Installer Software will allow for QC analysis of the data. The CPGR will assess the quality of the data firstly using a metric called the 'Detected CpG' Metric for each sample, which is an indication of the number of markers on the Bead Chip that were detected in that sample. Ideally 96% or more of the markers should be detected, however in Genome Studio this is not presented in a percentage and as there are 865918 CpG sites (markers) included on the bead chip, for a 96% call rate 831281 or more sites need to be detected. If a sample has fewer than 831281 markers detected, it will be deemed to have failed. Other metrics that can be assessed include the bisulphite conversion controls which are assessed through relative intensities of the probes in the red and green channels. BeadArray Controls Reporter Installer Software simplifies this interpretation and if used the thresholds specified in the User guide will be followed (file:///Z:/Service%20Projects/EPIC/beadarray-controls-reporter-user-guide 1000000004009-00.pdf).

The CPGR will present an analytical report at the end of each batch. The analytical report will include the data of the QC analysis done by the CPGR. Furthermore, the analytical report will include the workflow and experimental procedures, as well as any deviations. The results for all samples will be provided as IDAT files (raw data) either on a hard drive or via a secure portal for transferring the data as agreed upon by the client.

It is strongly advised to download the Methylation GenomeStudio Software 2011.1 software from Illumina using the following link: https://emea.support.illumina.com/softwaredownload.html/?assetId=a3cef505-e401-43ef a4d2-b265f800139d&assetDetails=genomestudio-v2011-1.zip. Various other software options are also available for secondary analysis of the methylation array results, please refer to the following website for tips regarding analysis methods: https://www.illumina.com/techniques/microarrays/methylation-arrays/methylation-array data-analysis-tips.html

**Managing sample number discrepancies**

In the event that the full number of samples are not delivered to the CPGR at the agreed upon date, the following provisions have been made:

The CPGR will allow an additional five working days for outstanding samples to be delivered.

**If at least 80% of the samples have been delivered by the stipulated time, the CPGR staff will engage the client to decide whether to go ahead with the project. If a go-ahead decision is agreed, no further samples will be accepted for the project, and costs associated with services not delivered will be forfeited.**

Where less than 80% of the sample number is delivered, or if the client is not willing to go ahead with the project using only 80% of the samples, the project will be re-scheduled according to instrument/staff availability and the delivery of outstanding samples. Purchased reagents will be kept at the CPGR in case the project is resumed. The client will be advised of the expiry date of the reagents so that the client is aware that project completion needs to be before the reagents expire.

If outstanding samples are not received within a 1-month time window, the project will be officially closed. At the discretion of CPGR management, unspent funds after reagent costs have been subtracted will either be returned to the client's account or a credit note will be passed.

The CPGR will charge an administration fee, and other fees as applicable, to the client in the event of a re-schedule or cancellation due to incomplete sample delivery (less than 80% of expected samples).

### Quality control and quality assurance

The MagMax DNA Multi-Sample Ultra 2.0 Kit typically yields 1.5 – 4 µg of DNA for every 50 µL of blood and the QIAamp DNA Mini typically yields 6 µg of DNA for 200 µl of whole blood. The QuantiFluor dsDNA quantification assay or Qubit BR DNA kit will be used to determine the dsDNA concentration of the extracted DNA sample. The Nanodrop spectrophotometer will be used to determine the DNA concentration as well as the presence of possible contaminants and/or inhibitors in the DNA. The integrity of samples will be assessed by performing gel electrophoresis on the samples using a 1-2% agarose gel with a suitable ladder to confirm the

presence of high molecular weight DNA. The samples need to conform to the requirements as listed under point 3.

Detected CpG sites need to exceed 96% for all samples depending on the quality of the DNA and the source from which it was obtained. Duplicate and control samples may be included upon the client's request to confirm concordance of replicates and monitor assay perforance, respectively.
