## Supplementary figures and images for "Wearable-ome meets epigenome: A novel approach to measuring biological age with wearable devices"

### supplementary image 1

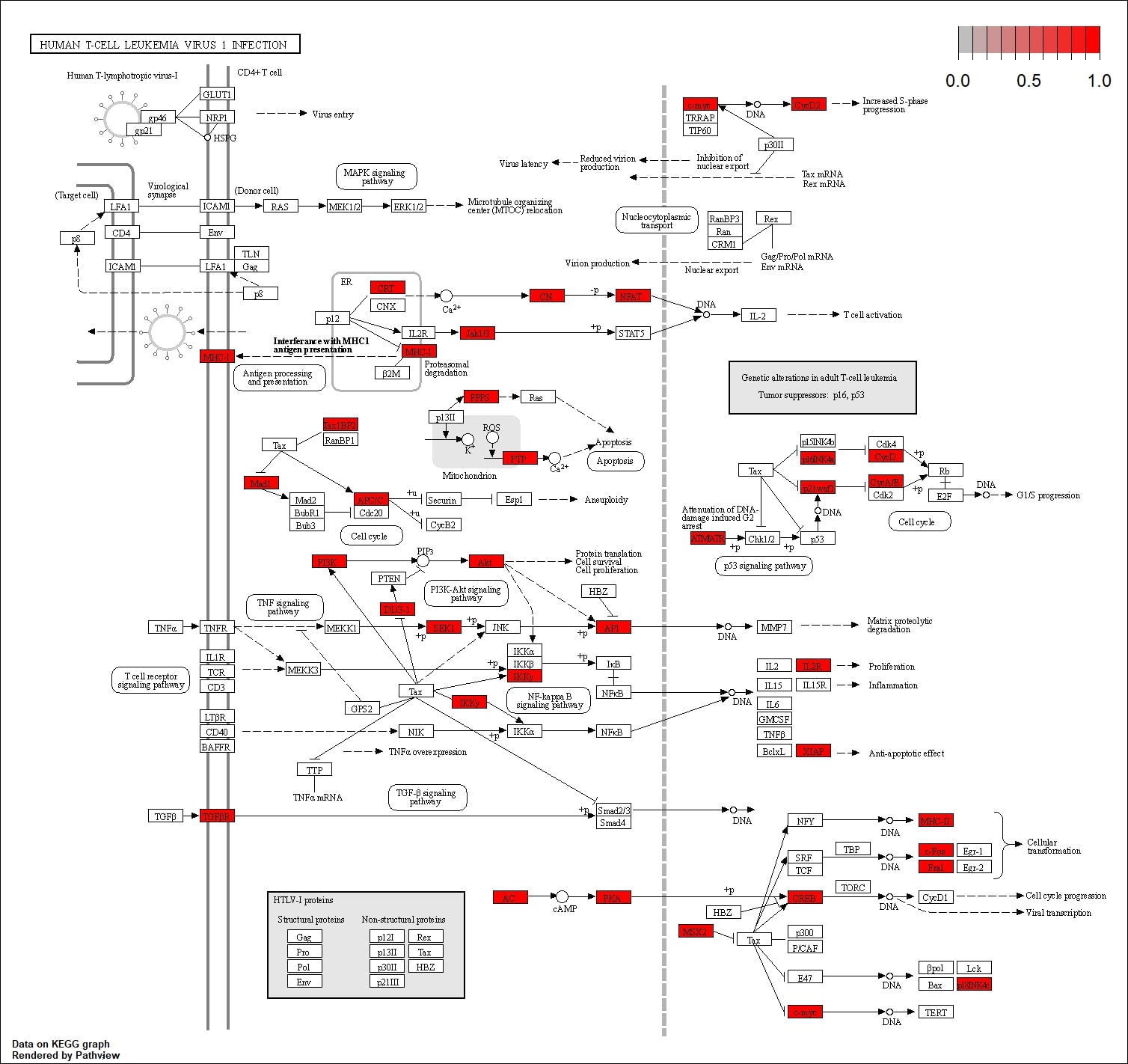
